# Tafazzin knockdown in murine mesenchymal stem cells enhances the tafazzin knockdown mediated elevation in interleukin-10 secretion from murine B lymphocytes

**DOI:** 10.1101/2023.03.01.530640

**Authors:** Hana M. Zegallai, Ejlal Abu-El-Rub, Grant M. Hatch

**Affiliations:** Department of Pharmacology & Therapeutics, University of Manitoba, Children’s Hospital Research Institute of Manitoba (CHRIM), Winnipeg, Manitoba, Canada; Physiology and Pathophysiology, Department of Basic Medical Sciences, Faculty of Medicine, Yarmouk University, Irbid, Jordan

**Keywords:** Barth Syndrome, tafazzin, cardiolipin, B lymphocytes, bone marrow mesenchymal stem cells, mitochondria, interleukin-10, immunosuppression, infection, cell proliferation, genetic diseases, lipopolysaccharide, neutropenia

## Abstract

Barth Syndrome is a rare X-linked genetic disorder caused by mutations in the *TAFAZZIN* gene. We recently demonstrated that tafazzin (Taz) protein deficiency in murine mesenchymal stems (MSCs) reduces immune function of activated wild type (WT) B lymphocytes. Interleukin-10 (IL-10) is a key anti-inflammatory cytokine capable of exerting immunosuppressive effects on myeloid cells. Here we examined if Taz deficiency in murine MSCs altered proliferation and IL-10 production in Taz deficient lipopolysaccharide (LPS)-activated murine B lymphocytes. Bone marrow MSCs and splenic B lymphocytes were isolated from WT or Taz knockdown (TazKD) mice. WT or Taz deficient MSCs were co-cultured with either LPS-activated WT or LPS-activated Taz deficient B lymphocytes for 24 h and B cell proliferation and IL-10 production determined. Taz deficient MSCs exhibited increased phosphatidylinositol-3-kinase (PI3K) mRNA expression compared to WT MSCs indicative of enhanced immunosuppression. Co-culture of Taz deficient MSCs with Taz deficient LPS-activated B cells resulted in a greater reduction in proliferation of B cells compared to Taz deficient MSCs co-cultured with LPS-activated WT B cells. In addition, co-culture of Taz deficient MSCs with Taz deficient LPS-activated B cells resulted in an enhanced production of IL-10 compared to Taz deficient MSCs co-cultured with LPS-activated WT B cells. Thus, Taz deficiency in murine MSCs potentiates the Taz knockdown-mediated elevation in IL-10 secretion from LPS-activated Taz knockdown B lymphocytes. These data suggest that Taz deficient MSCs may modulate the activity of other Taz deficient immune cells potentially promoting an enhanced immunosuppressive state.

## Introduction

Barth Syndrome (BTHS) is a rare X-linked genetic disease characterized by cardiomyopathy, skeletal myopathy, growth retardation, neutropenia, and 3-methylglutaconic aciduria^1,2^. It is caused by mutations in the *TAFAZZIN* gene localized to chromosome Xq28.12. The *TAFAZZIN* gene codes for the cardiolipin transacylase protein tafazzin (Taz).

Neutropenia is observed in many patients with BTHS and this may result in severe infections. Recurrent fever and infections in BTHS patients with neutropenia can be treated with granulocyte-colony stimulating factor (G-CSF) and antibiotics^3^. The molecular mechanisms underlying the neutropenia in BTHS are unknown. Interestingly, neutrophils isolated from BTHS patients exhibited normal motility, killing activity, phagocytosis, mitochondrial shape and mass^4^. A previous study indicated that in mice transplanted with TAFAZZIN-KO fetal liver cells, Taz deficiency did not affect hematopoiesis but resulted in leukopenia in the setting of normal peripheral red blood cell and platelet counts (Sohn et al 2022 Blood Adv.)5. The leukopenia manifested as a mild decrease across lineages, including peripheral B cells, T cells, myeloid cells, and neutrophils. Moreover, loss of Taz in granulocyte monocyte progenitors did not affect neutrophil phagocytosis, cytokine production, reactive oxygen species generation or mitochondrial complex assembly but increased sensitivity to endoplasmic reticulum stress in vivo.

There is limited information on the role of Taz deficiency on other cells of the immune system particularly in the context of cell-to-cell interaction of different immune cell types. We previously demonstrated that lipopolysaccharide (LPS)-activated B lymphocytes from Taz knockdown (TazKD) mice exhibited a reduced ability to secret cytokines, such as interleukin-10 (IL-10), which is known to attract neutrophils to sites of infection^6,7^. In addition, MSCs from TazKD mice impaired the function of LPS-activated wild type (WT) B lymphocytes. Thus, it is possible that altered immune cell-to-cell signaling in Taz deficient MSCs cells could impact the ability of other Taz-deficient immune cells to modulate neutrophil function and contribute to infections in BTHS.

In this study, we examined if Taz deficiency in murine MSCs altered proliferation and IL-10 production in Taz deficient LPS-activated murine B lymphocytes. We show that Taz deficiency in murine MSCs potentiates the elevation in IL-10 secretion from LPS-activated TazKD B lymphocytes when co-cultured together suggesting that Taz deficient MSCs may modulate the activity of other Taz deficient immune cells potentially promoting an enhanced immunosuppressive state.

## Materials and Methods

### Reagents

All reagents used, unless otherwise indicated, were of analytical grade and obtained from either Fisher Scientific (Winnipeg, MB) or Sigma-Aldrich (Burlington, MA).

### Animals

Experimental procedures were performed with approval of the University of Manitoba Animal Policy and Welfare Committee in accordance with the Canadian Council on Animal Care guidelines and regulation and are reported in accordance with ARRIVE guidelines. Animals were housed (12 h light/dark cycle) in a pathogen-free facility at the University of Manitoba. TazKD mice were generated by mating transgenic male mice containing doxycycline (dox) inducible Taz specific short-hair-pin RNA (shRNA) (B6. Cg-Gt(ROSA)26Sortm1(H1/tetO-RNAi:Taz,CAG-tetR) Bsf/ZkhuJwith female C57BL/6J mice from Jackson Laboratory (Boston, MA). Knock down was maintained postnatally by administering dox (625 mg of dox/kg of chow) as part of the rodent chow (Rodent diet TD.01306 from Harlan, Indianapolis, USA). WT and transgenic mice were maintained postnatally on the dox diet and mice aged 10-14 weeks were used for the experiments.

### Isolation and culture of cells

MSCs were isolated from the femurs and tibias of WT or TazKD mice by flushing the bone marrow cells and cultured as previously described^6^. Splenic naïve B lymphocytes from WT and TazKD mice were isolated and cultured as previously described^6^. For co-culture studies with WT or TazKD MSCs, naïve or TazKD B cells were first stimulated by incubation with LPS (10 μg/ml) for 24 h and were then co-cultured with MSCs for 24 h in 24-well plates at a ratio of 1:10 (MSCs:B cells) as previously descibed^6^.

### Measurement of PI3K mRNA expression, cell proliferation and IL-10 production

RNA was isolated from WT MSCs and TazKD MSCs using RNeasy Mini Kit and Qiashredder homogenizer columns according to the manufacturer’s instructions (Qiagen, ThermoFisher). RT-PCR for PI3K mRNA was performed using the QuantiTect® Probe RT PCR Kit and the double-stranded DNA stain SYBR green according to the manufacturer’s instructions (Qiagen, ThermoFisher). B-actin was used as the control. The primers used for RT-PCR detection are listed in **Table 1** and were obtained from Integrated DNA Technologies (Coralville, IA). Relative gene expression analysis was calculated using the 2-^ΔΔ^Ct method. B lymphocyte proliferation was measured using the cell proliferation assay kit (CellTiter 96 Aqueous) from Promega (Madison, WI) according to manufacturer’s instructions. IL-10 was measured in the supernatant using specific IL-10 antibody (Catalog no. sc-365858) from Santa Cruz Biotechnology (Dallas, TX) as previously described^6^.

**Table 1.**
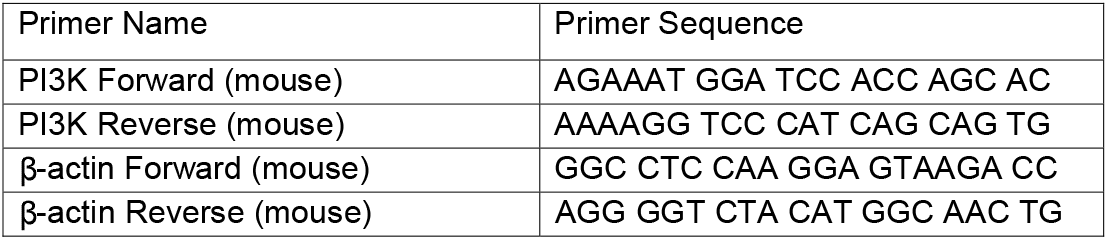
Primers used for RT PCR of PI3K.

### Statistical analysis

Data was expressed as mean ± standard deviation. Student’s t-test was used to compare between two groups. One-way analysis of variance followed by Tukey’s post-hoc multiple comparison test was performed to compare between multiple groups. A p value of <0.05 was defined as statistically significant.

## Results

Increased PI3K expression and activity is associated with enhancement of immunosuppression by MSCs^7^. We previously demonstrated that PI3K protein expression was elevated in MSCs isolated from TazKD mice^6^. MSCs were isolated from bone marrow of WT and TazKD mice and PI3K mRNA expression determined. PI3K mRNA expression was elevated in MSCs isolated from TazKD mice compared to WT controls (**Figure 1**). This confirmed the MSCs from TazKD mice used in this study exhibited an increased PI3K associated with enhanced immunosuppression^5,7^.

**Figure 1.**
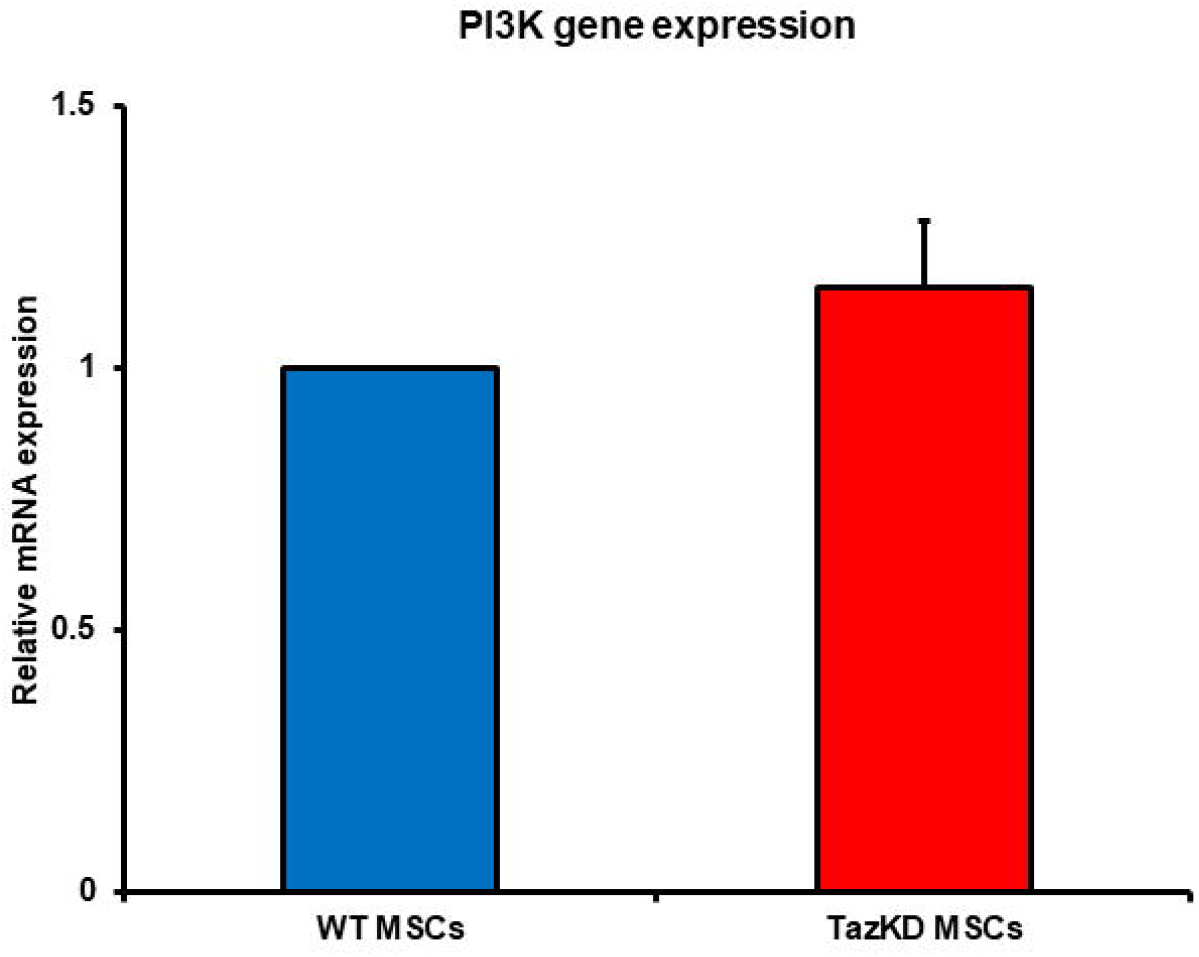
PI3K mRNA expression in TazKD MSCs. Bone marrow derived MSCs were isolated from WT or Taz knockdown mice and PI3K mRNA expression determined as described in Materials and Methods. Data represents the mean ± SD, n=4.

We previously demonstrated that TazKD MSCs impaired WT B lymphocyte proliferation^6^. To determine if TazKD MSCs impaired TazKD B lymphocyte proliferation, MSCs were isolated from bone marrow of WT and TazKD mice and co-cultured with either WT or TazKD splenic B lymphocytes for 24 and proliferation of B lymphocytes determined. As expected, the presence of either WT or TazKD MSCs significantly inhibited WT or TazKD B lymphocyte proliferation (**Figure 2**). Interestingly, the inhibition of B lymphocyte proliferation appeared greater when TazKD MSCs were co-cultured with TazKD B lymphocytes compared to TazKD MSCs co-cultured with WT B lymphocytes. Thus, deficiency of Taz in MSCs seemed to enhance the reduction in proliferation of Taz deficient B lymphocytes.

**Figure 2.**
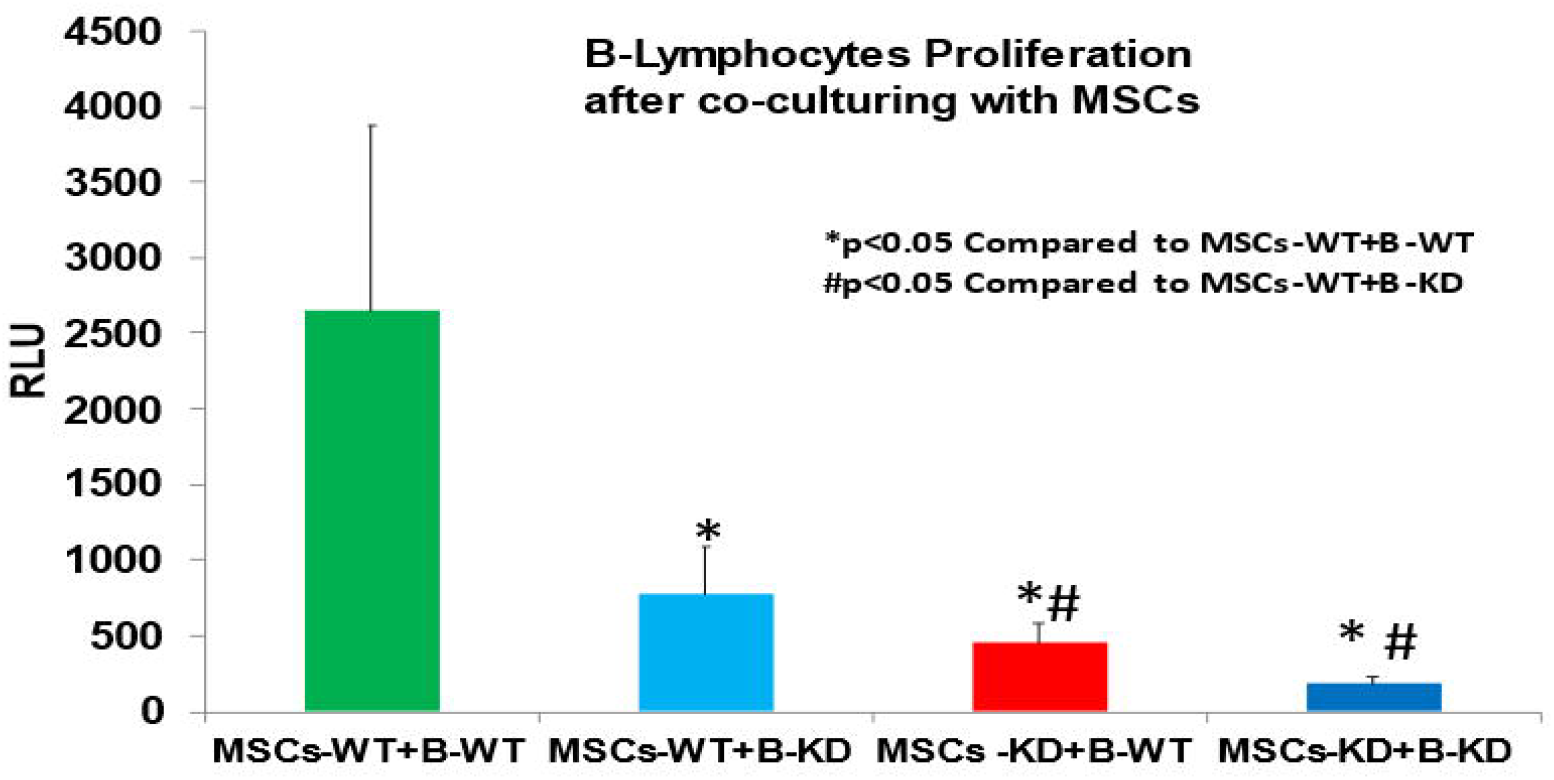
Proliferation of WT and TazKD B lymphocytes co-cultured with either WT or TazKD MSCs. Bone marrow MSCs and splenic B lymphocytes were isolated from WT or Taz knockdown mice. WT or Taz deficient MSCs were then co-cultured with either LPS-activated WT or LPS-activated Taz deficient B lymphocytes for 24 h and B cell proliferation determined as described in Materials and Methods. Data represents the mean ± SD, n=4. *p<0.05 compared to WT MSCs, #p<0.05 compared to TazKD MSCs.

IL-10 is a potent immunosuppressive cytokine^9^. We previously demonstrated that TazKD MSCs promoted enhanced IL-10 production in WT B lymphocyte^6^. To determine if TazKD MSCs impaired TazKD B lymphocyte IL-10 production, MSCs were isolated from bone marrow of WT and TazKD mice and co-cultured with either WT or TazKD splenic B lymphocytes for 24 and IL-10 secretion from lymphocytes determined. TazKD MSCs significantly increased IL-10 secretion from either WT or TazKD B lymphocytes (**Figure 3**). IL-10 production from TazKD B lymphocytes was significantly higher than that of WT B lymphocytes when co-cultured with TazKD MSCs. Thus, deficiency of Taz in MSCs enhanced IL-10 production in Taz deficient B lymphocytes.

**Figure 3.**
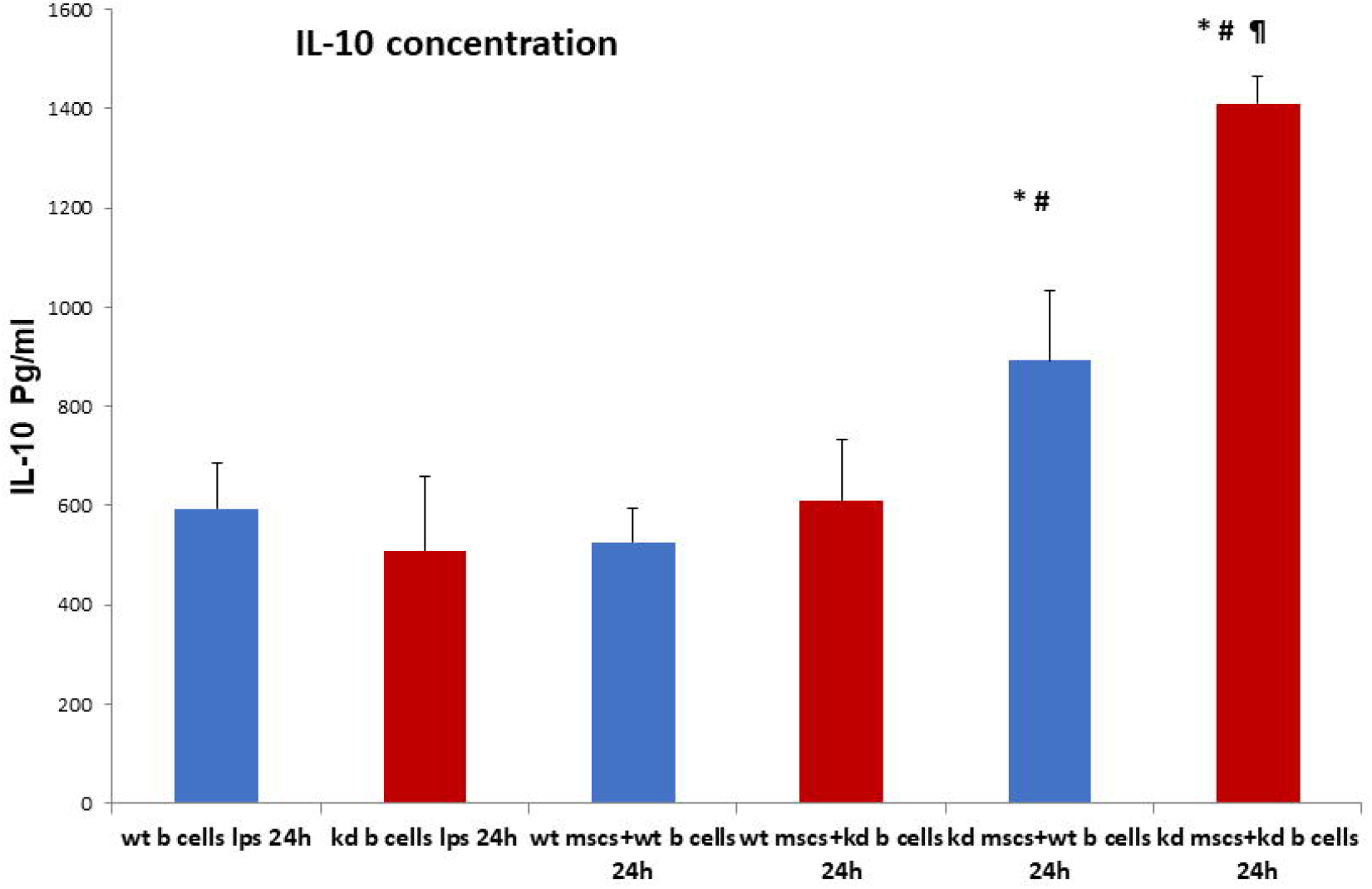
IL-10 secretion from WT and TazKD B lymphocytes co-cultured with either WT or TazKD MSCs. Bone marrow MSCs and splenic B lymphocytes were isolated from WT or Taz knockdown mice. WT or Taz deficient MSCs were then co-cultured with either LPS-activated WT or LPS-activated Taz deficient B lymphocytes for 24 h and IL-10 determined as described in Materials and Methods. Data represents the mean ± SD, n=4. *p<0.05 compared to WT B Cells, #p<0.05 compared to WT MSCs, ¶ p<0.05 compared to TazKD B cells.

## Discussion

The major findings of this study are 1. Taz deficient MSCs exhibit increased phosphatidylinositol-3-kinase (PI3K) mRNA expression compared to WT MSCs, 2. Co-culture of Taz deficient MSCs with Taz deficient LPS-activated B cells result in a greater reduction in proliferation of B cells compared to Taz deficient MSCs co-cultured with LPS-activated WT B cells, and 3. Co-culture of Taz deficient MSCs with Taz deficient LPS-activated B cells resulted in an enhanced production of IL-10 compared to Taz deficient MSCs co-cultured with LPS-activated WT B cells. Thus, Taz deficiency in murine MSCs may modulate the activity of other Taz deficient immune cells potentially promoting an enhanced immunosuppressive state.

Many BTHS boys exhibit neutropenia with increased susceptibility to infections^1,2^. However, the neutropenia does not explain why infection may occur despite consistent prevention of neutropenia in some BTHS patients. The clinical data, routine blood counts, and responses to granulocyte-colony stimulating factor therapy from 88 affected boys was recently reviewed^10^. It was concluded from that study that susceptibility to infections is due only in part to neutropenia since in some instances infections may occur despite consistent prevention of neutropenia through granulocyte-colony stimulating factor therapy. This is highlighted by the recent observation of a BTHS boy with hypogammaglobulinaemia and B cell lymphopaenia who required subcutaneous immunoglobulin replacement as adjunctive therapy with granulocyte-colony stimulating factor for infections^11^. In a previous study we observed elevated PI3K protein expression in TazKD MSCs compared to WT MSCs which is indicative of an enhanced immunosuppression^6^. In the current study, this was confirmed as Taz deficient MSCs exhibited increased phosphatidylinositol-3-kinase (PI3K) mRNA expression compared to WT MSCs. In addition, we previously demonstrated that TazKD B cells exhibited decreased cellular proliferation and increased IL-10 production compared to WT cells^7^. In the current study, we confirmed these observations and further observed that Taz deficient murine MSCs potentiated the Taz knockdown-mediated elevation in IL-10 secretion from LPS-activated TazKD B lymphocytes. These observations suggest that Taz deficient MSCs modulate the activity of other Taz deficient immune cells potentially promoting an enhanced immunosuppressive state. A limitation of our study is that the model we have used is not a complete knockdown as TazKD MSCs and B cells exhibit residual Taz protein^6,7^.

A recent study in which Taz knockout/X females in the C57BL6 inbred strain were mated with males from eight inbred strains indicated that genetic modifiers powerfully modulate the phenotypic expression of Taz loss-of-function^12^. These data may additionally, in part, explain why not all BTHS boys exhibit neutropenia. In summary, we hypothesize that attenuated immune system function in some BTHS patients may contribute to the presence of infections in these individuals even when neutrophil counts are at or near normal levels.

## Acknowledgements

We thank Marilyne Vandel for technical assistance and Dr. Laura K. Cole for preparation of the TazKD mice. H.M.Z. was supported by a Libyan North American Scholarship. This work was supported by a grant from the Natural Sciences and Engineering Research Council of Canada (NSERC) to G.M.H.

## Notes

### Competing Interest Statement

The authors have declared no competing interest.

### Summary of Updates

The revised text contains extra references

